# GCN5L1 inhibits pyruvate dehydrogenase phosphorylation during cardiac ischemia-reperfusion injury

**DOI:** 10.1101/2024.06.03.597148

**Authors:** Paramesha Bugga, Michael W Stoner, Janet R Manning, Bellina AS Mushala, Nisha Bhattarai, Maryam Sharifi-Sanjani, Iain Scott

**Author notes:** For correspondence: Iain Scott, PhD, FAHA, FCVS, University of Pittsburgh, BST W1048, 200 Lothrop Street, Pittsburgh, PA 15261.

## Abstract

Myocardial infarction remains one of the leading causes of mortality. Reperfusion of the infarcted myocardium restores blood flow and reduces primary ischemic injury. However, despite its protective function, reperfusion is also associated with several deleterious outcomes that can result in ischemia-reperfusion (I/R) injury to cardiac tissue. While negative outcomes such as reactive oxygen species generation are strongly associated with I/R injury, cardiac energy metabolism is also greatly disrupted. Furthermore, previous studies have shown that the restoration of normal fuel oxidation in the myocardium regulates the extent of contractile recovery. A better understanding of the pathophysiological mechanisms underlying I/R injury may allow us to develop new treatments that limit the negative aspects of the process. In this study, we examined the role played by GCN5L1, a protein implicated in the regulation of energy metabolism, in I/R injury. We demonstrate that cardiac-specific loss of GCN5L1 promotes the inhibitory phosphorylation of pyruvate dehydrogenase *in vitro* and *in vivo*, a process likely to inhibit glucose oxidation, and that this corresponds to increased myocardial damage following ischemia-reperfusion (I/R) injury.

## INTRODUCTION

Ischemic heart disease is the leading cause of death among all cardiovascular diseases (GDB 2021 Global Causes of Death Collaborators, 2024). In ischemic heart disease conditions, cardiomyocytes are deprived of oxygen and nutrients, which leads to cardiomyocyte death and ultimately heart failure. This process is most acutely evident in myocardial infarction, where the complete occlusion of coronary arteries results in a loss of perfusion of the myocardium, resulting in cardiomyocyte apoptosis and eventual scar formation (reviewed in Shi et al, 2025). To reduce the extent of tissue damage, affected individuals typically undergo treatment with thrombolytic drugs, or receive invasive procedures such as percutaneous coronary intervention to reopen occluded arteries (Shi et al, 2025). While these treatments restore blood flow to the myocardium and protect against further primary ischemic injury, the reperfusion process can also lead to secondary ischemia-reperfusion (I/R) injury. This occurs through several mechanisms, including excessive reactive oxygen species production, mitochondrial dysfunction, and the activation of apoptotic and inflammatory signaling pathways (Shi et al, 2025).

Beyond these well-described aspects of treatment for myocardial infarction, there are numerous changes in cardiac energy metabolism that play an important role. The healthy adult heart typically obtains most of its energy for contraction through the oxidation of fatty acids, with the remainder coming from glucose/pyruvate oxidation, ketone oxidation, and glycolysis (Lopaschuk and Ussher, 2016). During ischemic injury, the loss of oxygen supply severely limits cardiac energy production, with the non-oxidative glycolysis process being the most important (Lopaschuk et al, 2021). The return to normal oxidative fuel metabolism is a major factor in the restoration of contractile function, and may regulate the extent of recovery (Harmancey et al, 2013; Kolwicz et al, 2015). One key aspect of this process is the recoupling of glycolysis to glucose/pyruvate oxidation as oxygen supply returns, and delays in this process can inhibit full recovery (Masoud et al, 2014). Finally, in terms of ATP produced per mole of O_2_ consumed, glucose is a more efficient fuel than fatty acids and ketones (Lopaschuk and Ussher, 2016), which may be particularly important under reduced oxygen conditions. As such, a better understanding of the pathways that regulate energy metabolism in the heart, particularly in terms of glucose handling, may permit the development of new therapeutic strategies to limit disease.

We previously demonstrated that GCN5L1, a mitochondrial- and cytosolic-localized protein, regulates fuel metabolism in cardiac cells (Thapa et al, 2017; Manning et al, 2019a). Using an *in vitro* model, this previous work showed that the genetic downregulation of GCN5L1 abundance in cardiac cells resulted in decreased glucose oxidation, and an increased reliance on glycolysis, under normoxic conditions through an unknown mechanism (Manning et al, 2019a). Here, we show that genetic downregulation of GCN5L1 *in vitro*, and cardiac-specific deletion *in vivo*, promotes the inhibitory phosphorylation of pyruvate dehydrogenase (PDH), the key regulator of glucose/pyruvate oxidation. This occurs in concert with increased tissue damage in cardiac-specific GCN5L1 knockout mice after transient ischemia/reperfusion injury surgery, suggesting that GCN5L1 expression in the heart protects against the maladaptive loss of normal glucose oxidation in ischemic injury.

## MATERIALS AND METHODS

### Generation of cardiac-specific GCN5L1 knockout mice

All mice were bred and housed according to the University of Pittsburgh’s IACUC protocols. At our request, EUCOMM created cardiomyocyte-specific conditional knockouts on a C57BL/6J background by crossing αMHC-Cre (Jax B6.FVB (129)-Tg (Myh6-cre/Esr1*)1JMK/J) mice (i.e. αMHC-MerCreMer) to mice with LoxP sites introduced around exon 3 of BLOC1S1 (GCN5L1^FL^). Cardiomyocyte-restricted knockout mice were produced by a single 40 mg/kg i.p. tamoxifen injection and confirmed with qPCR and western blotting 4 weeks post-tamoxifen injection.

### Cardiac ischemia-reperfusion injury surgery model

For transient coronary artery ligation (CAL) surgery, male mice were anesthetized with 3% isoflurane in an induction chamber before intubation. Dorsal surface hair was removed by applying depilatory cream, thoracotomy performed between the 3rd and 4th intercoastal ribs, and the heart exposed by using homemade retractors. The left anterior descending coronary artery was identified below the left atrium (∼2mm), and occluded with an 8-0 suture knot for 45 min. Reperfusion was performed by removing the occlusion, and allowed to proceed for 24 h. After reperfusion was initiated, the intercoastal muscles and ribs were closed with 6-0 silk sutures, and the skin closed with 6-0 silk sutures in an interrupted pattern. Cardiac function and myocardial strain were measured after 24 h of reperfusion. For sham operated groups, we followed a similar procedure except for coronary artery occlusion.

### Echocardiography

To evaluate the cardiac function and structure, we performed transthoracic echocardiography using a VisualSonics Vevo 3100 system (Visual Sonics, Toronto, Ontario, Canada) with an MS400 (30 MHz centerline frequency) probe. Mice were anesthetized with isoflurane (induction 3% and maintenance 1.5–2%), and hair was removed with depilatory cream. Mice were placed in a supine position on a heated table (core temperature maintained at 37°C) with embedded ECG leads. B- and M-mode images were acquired from a parasternal short-axis view to evaluate left ventricle (LV) ejection fraction (EF), fractional shortening (FS), and volume at systole/diastole by M-mode images.

### Cardiac Markers in Ischemic-reperfusion Injury

Serum lactate dehydrogenase (LDH) and troponin, markers of cardiomyocyte damage following ischemic injury, were measured using commercially available kits.

### Cell culture

AC16 cells were purchased from Sigma-Millipore, and were grown under standard conditions (20% O_2_/5% CO_2_ at 37 ^0^C, in the presence of 25 mM glucose DMEM, 10% FBS, and antibiotic-antimycotic.

### Generation of GCN5L1 KD and overexpressed stable cell lines

AC16 cells were transduced with scrambled control or GCN5L1 shRNA lentiviral particles (Sigma-Aldrich, USA), or control or GCN5L1 ORF lentiviral particles (Origene, USA), at a multiplicity of infection (MOI) of 10. Cells were selected by puromycin. GCN5L1 knockdown (KD) or overexpression (OE) was confirmed by RT-qPCR and western blotting.

### Hypoxia/reoxygenation experiments

After 24 h of growth in 10 cm cell culture plates, hypoxia studies were performed on control and GCN5L1 KD AC16 cells. Cells were washed three times with 1X PBS, then hypoxic cells incubated with Esumi Buffer (12 mM KCl, 0.9 mM CaCl_2_, 0.5 mM MgCl_2_, 137 mM NaCl, 20 mM HEPES, 20 mM 2-deoxy-D-glucose (2-DG), and 20 mM actic acid, pH 6.2) for 3 h, followed by 1 h reoxygenation. After completion of hypoxia/reoxygenation treatment, cells were washed with 1X PBS and lysed in CHAPS lysis buffer with protease, phosphatase, and deacetylase inhibitors.

### Immunoblotting

Cells and cardiac tissues were lysed in 1% CHAPS lysis buffer. Protein samples were quantified using a μLITE analyzer (BioDrop, USA) and 25 μg of proteins were separated on an SDS-PAGE gel. Proteins were transferred onto nitrocellulose membrane and blocked using Odyssey blocking buffer (Li-Cor, USA). Membranes were incubated with primary antibodies (GAPDH, Cell Signaling, Cat. 97166 [D4C6R], mouse mAb, 1:1000; (αTubulin, Cell Signaling, Cat. 3873 [DM1A], mouse mAb, 1:1000; PDH, Cell Signaling, Cat. 3205S [C54G1], rabbit mAb, 1:1000; p-PDH[Ser293], Novo-Bioscience, Cat. NB110-93477, PDK4, Thermo Scientific, Cat. PA5-102685, rabbit mAb 1:1000; GCN5L1 1:500, rabbit pAb, validated in Bugga et al, 2024) overnight at 4 ^0^C. Membranes were washed with PBS-T, and developed with secondary antibodies followed by incubation at room temperature with fluorescent secondary antibodies for 1 h (800 nm anti-rabbit, 700 nm anti-mouse) at room temperature. Protein bands were visualized by an Odyssey Imager and analyzed using Image-J software.

### Statistical analysis

Statistical analyses were performed with GraphPad Prism. One-way ANOVA followed by Tukey’s multiple comparison tests was used for more than two experimental groups. A two-tailed student’s t-test was used for comparisons between two groups. A *P* value < 0.05 was considered statistically significant. Data are shown as Mean ± SEM.

## RESULTS

### GCN5L1 abundance is inversely correlated with pyruvate dehydrogenase phosphorylation in cardiac AC16 cells

We previously demonstrated that loss of GCN5L1 expression in cardiac cells led to an increased reliance on glycolysis for energy production under aerobic conditions (Manning et al, 2019a). This change occurred without a decrease in glucose uptake, suggesting that GCN5L1 protein abundance regulates the fate of glucose through an unknown mechanism. To better understand the pathways involved, we modulated GCN5L1 abundance via stable overexpression (OE) or knockdown (KD) in human cardiac AC16 cells. GCN5L1 OE led to a ∼50% decrease in inhibitory PDH phosphorylation (*P =* 0.06), while GCN5L1 KD led to a significant increase in PDH phosphorylation of the same magnitude (**Fig. 1A-D**). The increase in PDH phosphorylation in GCN5L1 KD cells was likely mediated by an increase in the abundance of the PDH kinase, PDK4, which phosphorylates PDH at Ser-293 (**Fig. 1E**). We observed no compensatory increase in the abundance of the PDH phosphatase, PDP1, and the increased PDH phosphorylation was likely compounded by an increase in the PDP1 regulatory subunit, PDPR, which inhibits PDP1 activity (Yan et al, 1996). Combined, these results suggest that GCN5L1 protein abundance in cardiac cells is inversely related to PDH phosphorylation status and activity.

**Figure 1:**
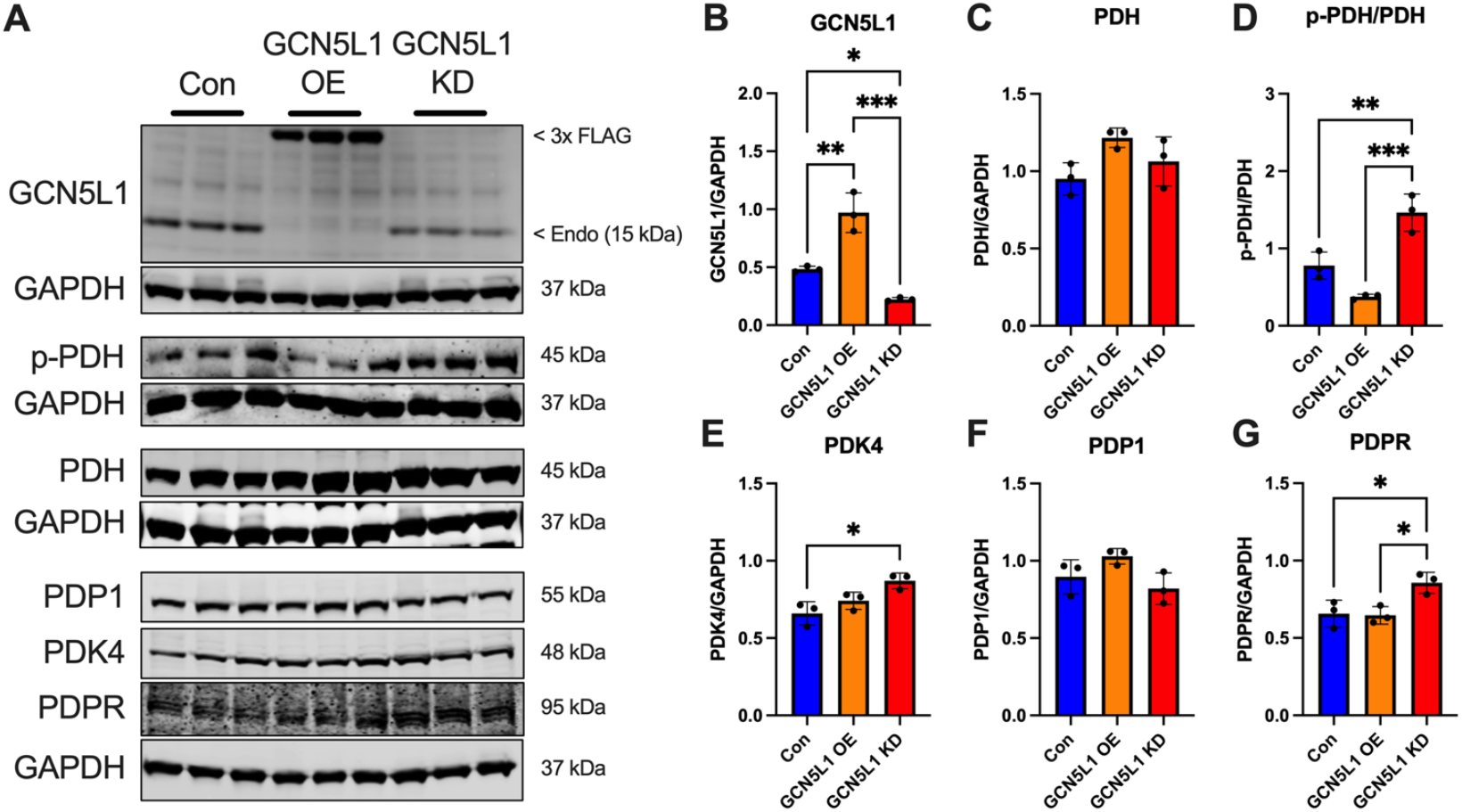
GCN5L1 abundance is inversely correlated with pyruvate dehydrogenase phosphorylation in cardiac AC16 cells. A. Representative immunoblots from stable cardiac AC16 cell lines with overexpression (OE) or knockdown (KD) of GCN5L1. In cells overexpressing GCN5L1, the endogenous protein is largely absent, suggesting a homeostatic response to limit total GCN5L1 levels. **B-G**. Protein abundance of GCN5L1, PDH, p-PDH, PDK4, PDP1, and PDPR in GCN5L1 OE or KD cells. N = 3; * = *P* < 0.05, ** = *P* < 0.01, *** = *P* < 0.001; one-way ANOVA with Tukey’s post-hoc test.

### GCN5L1 knockdown in AC16 cells leads to increased abundance of pyruvate dehydrogenase inhibitory proteins after hypoxia/reoxygenation stress

Upon entering an ischemic/hypoxic state, PDH becomes phosphorylated, which supports a significant decrease in glucose/pyruvate oxidation activity (Browning et al, 1981). We therefore tested whether GCN5L1 downregulation, which leads to PDH inhibitory phosphorylation in normoxia (**Fig. 1**), would have an impact on the levels of PDH regulatory proteins in hypoxia. PDH phosphorylation levels were significantly increased in both control and GCN5L1 KD AC16 cells following hypoxia/reoxygenation (H/R). However, there was no difference between the levels of p-PDH between the control and GCN5L1 KD H/R groups, suggesting that a maximal level of PDH phosphorylation had been reached following this treatment that was not GCN5L1-dependent (**Fig. 2A-D**). Despite this, we did detect a significant increase in both PDK4 and PDPR abundance in GCN5L1 KD H/R cells relative to the control group (**Fig. 2E,G**). However, this increase (which would have normally led to increased PDH phosphorylation) may have been partially negated by a non-significant ∼30% increase in PDP1 protein expression (**Fig. 2F**). Combined, these data suggest that loss of GCN5L1 expression in cardiac cells drives expression of proteins that inhibit PDH in both normoxia (**Fig. 1**) and hypoxia (**Fig. 2**).

**Figure 2:**
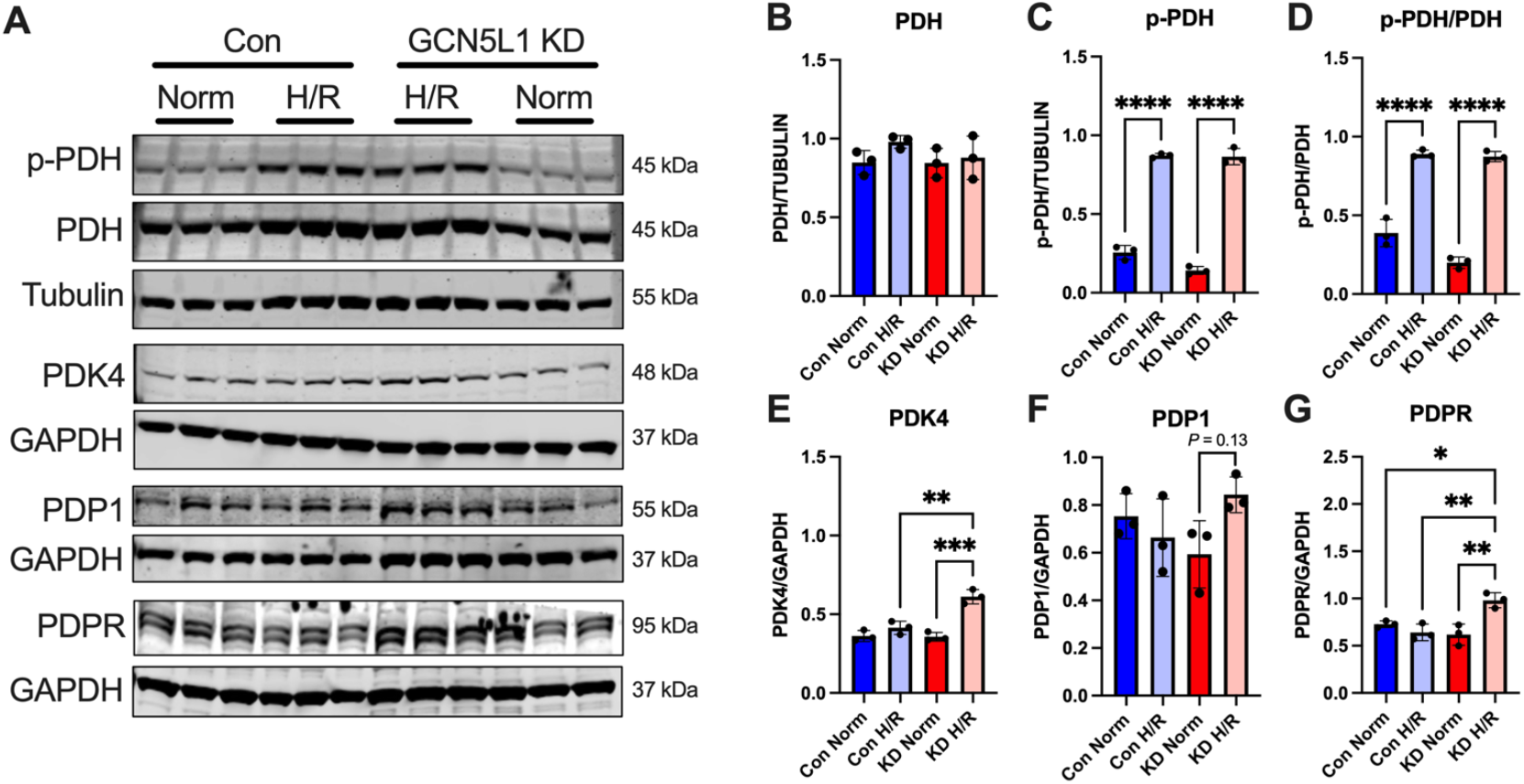
GCN5L1 knockdown in AC16 cells leads to increased abundance of pyruvate dehydrogenase inhibitory proteins after hypoxia/reoxygenation stress. A. Representative immunoblots from control or GCN5L1 KD AC16 cells after normoxia (Norm) or hypoxia/reoxygenation (H/R) treatment. **B-G**. Protein abundance of PDH, p-PDH, PDK4, PDP1, and PDPR in control and GCN5L1 KD AC16 cells after normoxia or hypoxia/reoxygenation. N = 3; * = *P* < 0.05, ** = *P* < 0.01, *** = *P* < 0.001, **** = *P* < 0.0001; one-way ANOVA with Tukey’s post-hoc test.

### Inhibitory pyruvate dehydrogenase phosphorylation is elevated in GCN5L1 cKO hearts following ischemia-reperfusion injury

Our previous studies on GCN5L1 in hypoxic conditions focused on the use of either cell or *ex vivo* tissue models (Manning et al, 2019a; Manning et al, 2019b). To determine whether GCN5L1 deletion in cardiomyocytes contributes to the response to hypoxia *in vivo*, we examined PDH phosphorylation in inducible, cardiac-specific GCN5L1 knockout (GCN5L1 cKO mice; Manning et al, 2019a). Wildtype (WT) and GCN5L1 cKO mice were subjected to transient ischemia via surgical constriction of the left descending coronary artery for 45 min, followed by 24 h of reperfusion. There was a significant increase in GCN5L1 expression in WT mice after ischemia-reperfusion (I/R) injury, which was absent in GCN5L1 cKO mice (**Fig. 3A,B**). When examining PDH phosphorylation, we found that there was a non-significant increase in p-PDH in GCN5L1 cKO mice under normoxia, which became a significant increase after I/R injury. Combined, these data suggest that GCN5L1 may aid the PDH dephosphorylation process after ischemia, which helps to recouple glycolysis to glucose/pyruvate oxidation.

**Figure 3:**
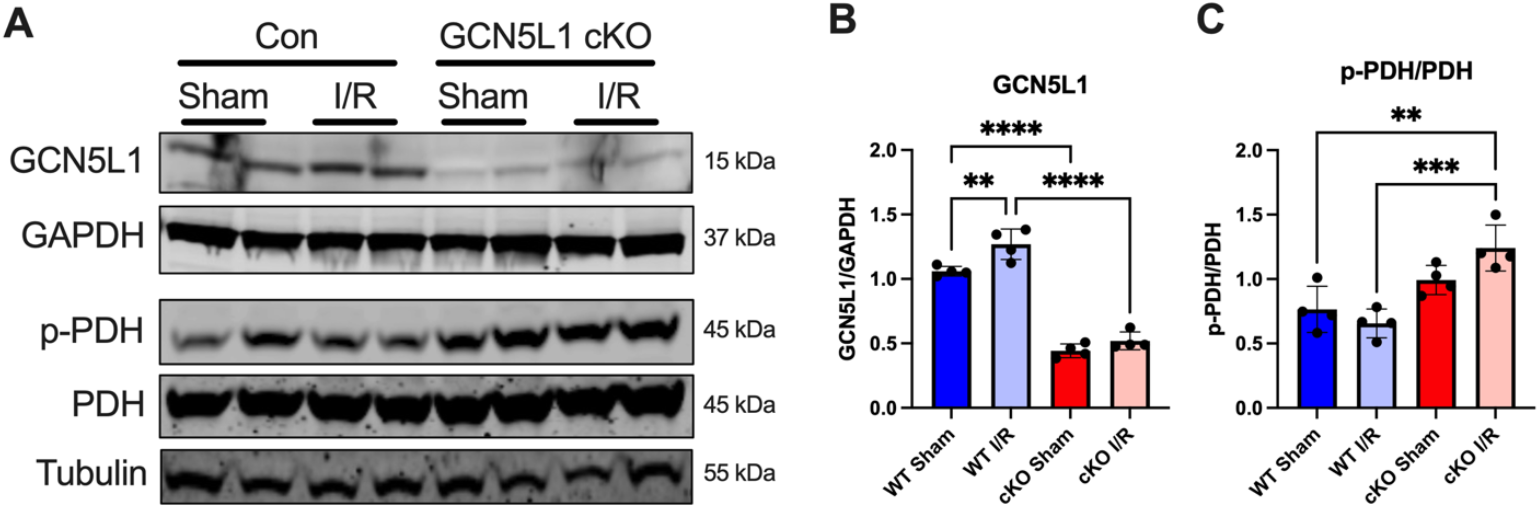
Inhibitory pyruvate dehydrogenase phosphorylation is elevated in GCN5L1 cKO hearts following ischemia-reperfusion injury. A. Representative immunoblots from wildtype (Con) and cardiac-specific GCN5L1 knockout (GCN5L1 cKO) mouse hearts after sham or ischemia-reperfusion (I/R) injury surgery. **B-C**. Protein abundance of GCN5L1 and p-PDH in wildtype or GCN5L1 cKO mouse hearts after sham or ischemia/reperfusion injury surgery. N = 4; ** = *P* < 0.01, *** = *P* < 0.001, **** = *P* < 0.0001; one-way ANOVA with Tukey’s post-hoc test.

### Loss of cardiomyocyte GCN5L1 expression leads to elevated tissue damage markers in the absence of functional decline

Finally, we examined whether the observed increase in PDH phosphorylation after ischemia in GCN5L1 cKO mice had functional or pathophysiological consequences. First, we assessed systolic function in WT and GCN5L1 cKO mice 24 h after reperfusion using echocardiography. While ejection fraction, fractional shortening, and left ventricular systolic volume were all significantly changed following I/R injury, there was no significant differences between the two genotypes (**Fig. 4A-D**). Secondly, we assessed whether markers of cardiac tissue damage were affected by GCN5L1 loss after I/R injury. We found that serum levels of cardiac troponin and lactate dehydrogenase were significantly increased in GCN5L1 cKO mice after ischemic injury relative to WT animals under the same conditions (**Fig. 4E,F**). Combined, these data suggest that loss of GCN5L1 in the myocardium exacerbates cardiac tissue injury following ischemia in the absence of early differences in contractile function.

**Figure 4:**
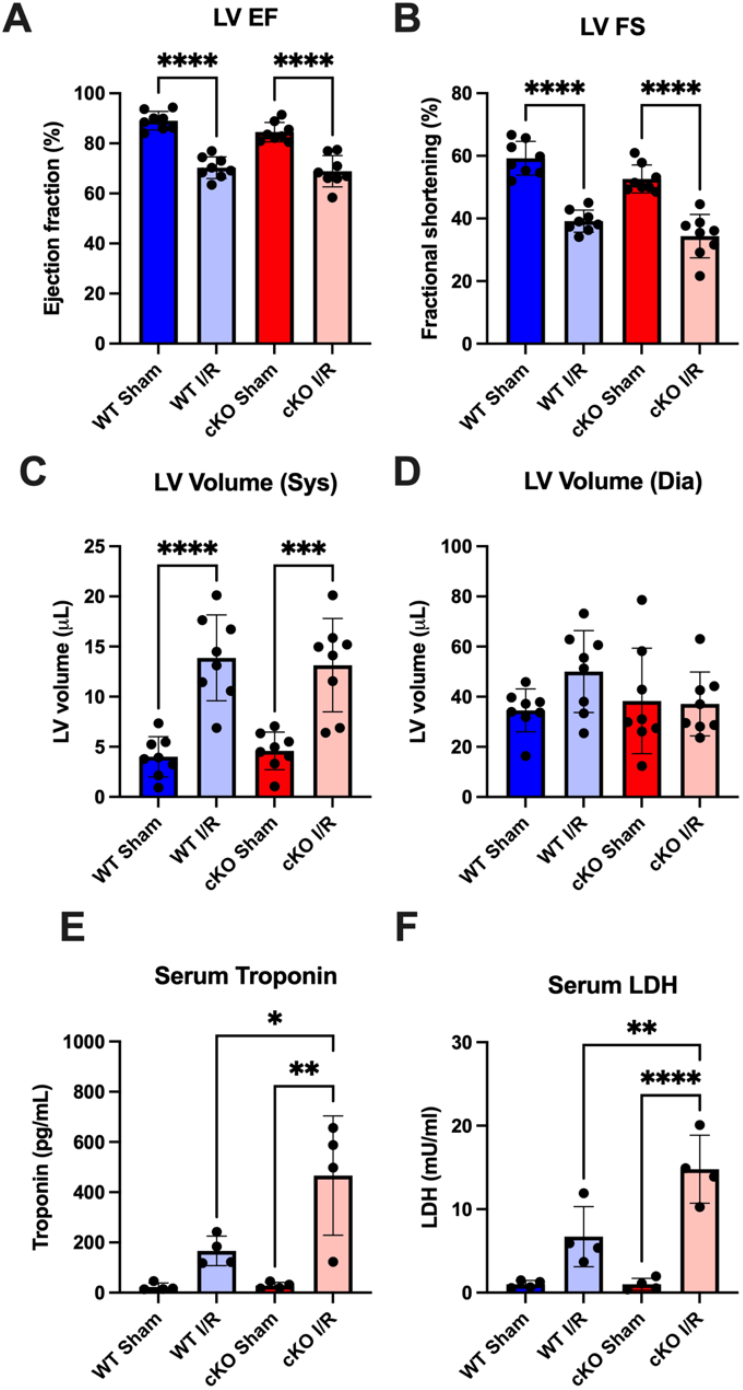
Loss of cardiomyocyte GCN5L1 expression leads to elevated tissue damage markers in the absence of functional decline. A-D. Echocardiographic analysis of left ventricle (LV) systolic function (EF, ejection fraction; FS, fractional shortening) and chamber volume at systole (Sys) and diastole (Dia) in wildtype and GCN5L1 cKO mice 24 h after sham or I/R surgery. N = 7-8. **E-F**. Serum troponin and lactate dehydrogenase (LDH) 24 h after sham or I/R surgery in wildtype and GCN5L1 cKO mice. N = 4; * = *P* < 0.05, ** = *P* < 0.01, *** = *P* < 0.001, **** = *P* < 0.0001; one-way ANOVA with Tukey’s post-hoc test.

## DISCUSSION

Changes in fuel metabolism are an inherent component of the response to ischemia in the heart (Lopaschuk et al, 2021). The switch from oxidative to glycolytic glucose utilization – necessitated by the loss of available oxygen – leads to reduced ATP generation and a deleterious change in ionic balance in the myocardium (Fillmore et al, 2014). While fatty acid oxidation recommences relatively quickly after reperfusion, the restoration of glucose/pyruvate oxidation is often delayed, and this is linked to a slower rate of functional recovery (Ussher et al, 2012). Better understanding the mechanisms that regulate the restoration of oxidative glucose use after ischemia may therefore help us to identify new therapeutic targets to promote recovery after myocardial infarction.

In this study, we show that GCN5L1 – a known regulator of cardiac fuel metabolism (Thapa et al, 2017; Manning et al, 2019a) – plays a key regulatory role in the activity of the PDH complex. We have previously shown that under non-ischemic, nutrient-excess conditions, GCN5L1 can acetylate and inhibit PDH activity (Thapa et al, 2022). In contrast, we show here that under normal nutrient and/or ischemic conditions, GCN5L1 expression is required to inhibit inhibitory phosphorylation of PDH by PDK4 (**Figs. 1**,**2**). Genetic depletion of GCN5L1 in cardiac cells results increased PDK4 gene expression (data not shown), and subsequent increases in PDK4 protein abundance (**Fig. 1**). This results in hyperphosporylation of the PDH complex, which inhibits its enzymatic activity (Browning et al, 1981). Loss of GCN5L1 expression promotes an additional increase in PDK4 expression under ischemic conditions *in vitro*, and a subsequent increase in PDH phosphorylation after I/R injury *in vivo* (**Figs. 2**,**3**). The increase in PDH phosphorylation in GCN5L1 cKO mice after I/R injury is subsequently linked to increased cardiac tissue damage (**Fig. 4**), suggesting that GCN5L1 is required to protect the heart from ischemic damage.

While loss of GCN5L1 in cardiomyocytes led to increased tissue damage after I/R injury (**Fig. 4E,F**), we did not observe any difference in cardiac functional activity or structural remodeling under the same conditions (**Fig. 4A-D**). One of the limitations of this study is that we measured cardiac function and recovered organs 24 h after the I/R injury surgery was performed, which did not allow us to examine longer-term functional recovery via compensatory structural remodeling. Future studies in this regard, where recovery after the I/R injury would be allowed to progress for 2-4 weeks, would allow us to determine if the increased initial tissue damage observed in GCN5L1 cKO mice (**Fig. 4E,F**) results in functional decline relative to WT mice in the long term.

Our study demonstrated that PDK4 was regulated by GCN5L1 expression, and that there was an inverse relationship between GCN5L1 and PDK4 abunance (**Fig. 1**). However, the mechanism underlying this regulation remains to be determined. Loss of GCN5L1 in the liver has been linked with changes in FoxO1 expression (Wang et al, 2017), which has been shown in the heart to regulate PDK4 expression (Gopal et al, 2017). Further studies will be required to determine the direct relationship between GCN5L1 abundance and PDK4 expression. In a similar vein, further work will be required to understand how GCN5L1 helps to regulate the expression of PDPR, the regulatory subunit of the PDH phosphatase protein PDP1. There have been few studies on PDPR, with early work showing that it negatively regulates PDP1 by inhibiting its Mg2^+^-dependent catalytic activity (Yan et al, 1996). Our results show that increased PDPR expression may work in concert with increased PDK4 expression (**Figs. 1**,**2**) to increase PDH phosphorylation. However, whether there is specific co-regulation of these two proteins remains to be determined.

In summary, we show that GCN5L1 protein abundance in the heart is negatively correlated with PDK4 expression, and that genetic depletion of GCN5L1 leads to increased inhibitory PDH phosphorylation and cardiac tissue damage in ischemia. These findings suggest that GCN5L1 may be an important component of post-ischemic restoration of cardiac glucose/pyruvate oxidation, and subsequent functional recovery.

## AUTHOR CONTRIBUTIONS

P.B. and I.S. conceived the study and designed experiments. P.B. performed experiments. P.B. analyzed data. P.B. and I.S. prepared figures. M.W.S., J.R.M., B.A.S.M., N.B., and M.S-S. provided critical input and expertise. P.B. and I.S. drafted the manuscript. P.B. and I.S. edited and revised the manuscript. All authors approved the final submission.

## ACKNOWLEDGEMENTS

This work was supported by National Institute of Health Fellowships (F31DK134089 and T32HL110849) to B.A.S.M, and National Institute of Health Research Grants (R01HL147861, R0HL156874) and an American Heart Association Established Investigator Award (23EIA1037834) to I.S. Echocardiography was carried out by the University of Pittsburgh Rodent Ultrasonography Core, which received funding from the NIH Shared Instrumentation Grant Program (S10OD023684).

